# Circadian Oscillation Detection Analysis and Comparison (CODAC): a Multicriteria Method to Estimate and Compare Rhythmicity

**DOI:** 10.64898/2026.08.17.745071

**Authors:** Thiago Parente da Silveira, Karla Lincoln, Thomas Nguyen, Leonardo Vinicius Monteiro de Assis

**Affiliations:** Department of Mathematics, Aeronautics Institute of Technology, São José dos Campos, SP, Brazil; Department of Chemistry and Molecular Biology, University of Gothenburg, Gothenburg, Sweden; Institute of Neurobiology, Center of Brain, Behavior and Metabolism, University of Lübeck, Lübeck, Germany; Wallenberg Centre for Molecular and Translational Medicine, University of Gothenburg, Gothenburg, Sweden; University Hospital Schleswig-Holstein, Campus Lübeck, Lübeck, Germany

**Author notes:** **Corresponding authors:** Leonardo Vinicius Monteiro de Assis | Thiago Parente da Silveira, | r.

**Keywords:** circadian rhythm, nonlinear optimization, rhythmicity, time-series analysis, omics, circadian bioinformatics

## Abstract

Analysis of circadian patterns in time-series data requires computational methods that can accommodate several factors, including variable sampling resolution, replicate number, and missing values. Most existing tools simplify rhythmicity to a strict dichotomy based solely on a single p-value threshold. This leads to a level of uncertainty that affects many biological targets. We developed CODAC (Circadian Oscillation Detection Analysis and Comparison), a framework that integrates nonlinear constrained optimization with a multicriteria rhythmicity classification scheme to evaluate rhythmic patterns without relying on a single statistical cutoff. This approach allows CODAC to identify and exclude medium-confidence rhythms rather than force them into a rhythmic/arrhythmic dichotomy. CODAC comprises four modules: (i) CODAC_single estimates rhythmicity within a single group; (ii) CODAC_flex extends this to identify distinct waveform types within one group; (iii) CODAC_compare performs pairwise comparisons across two or more groups to detect rhythmic or arrhythmic changes; and (iv) CODAC_multi handles more complex designs involving multiple-group comparisons. Using *in silico* simulations and public transcriptomic datasets, we show that CODAC performs comparably to established methods while providing additional flexibility for rhythm classification and comparison. Taken together, CODAC provides a flexible and open-source package for circadian timeseries analysis with automated visualization tools.

## 1 Introduction

Circadian rhythms are a fundamental property of life, allowing organisms to anticipate and adapt to predictable daily changes in the environment. In mammals, these rhythms emerge from cell-autonomous molecular clocks that coordinate biological processes across tissues and organs [1]. At the cellular level, circadian regulation extends across multiple layers of molecular organization, including rhythmic chromatin remodeling, transcription factor occupancy, RNA polymerase recruitment, nascent transcription, mRNA abundance, translation, and posttranslational modifications [2, 3, 4, 5, 6]. The expansion of high-throughput omics technologies has therefore transformed circadian biology, enabling rhythmicity to be investigated across different molecular layers, tissues, and experimental conditions. However, the omics revolution has also increased the need for robust computational methods capable of estimating rhythmicity across different platforms (e.g., RNA-seq and proteomics) and experimental conditions.

Computational methods for analyzing rhythmicity in biological time-series data can be broadly classified as non-parametric or parametric according to the assumptions they make about the underlying oscillatory pattern. Non-parametric, rank-based methods such as JTK_CYCLE and RAIN detect rhythmicity by evaluating the temporal ordering of observations rather than fitting a continuous mathematical function to the data [7, 8]. In contrast, parametric and model-based approaches explicitly represent rhythmicity using mathematical functions. Sinusoidal approaches, including classical cosinor analysis, typically fit a cosine function, or an equivalent combination of sine and cosine terms, to estimate the Midline-Estimating Statiscs of Rhythm (MESOR), amplitude, and phase, with the period either prespecified or estimated. Regression-based extensions such as CircaCompare, DryR, CompareRhythms, and diffCircadian use related model-based frameworks to detect rhythmicity and test differences in rhythmic parameters or overall rhythmic patterns between experimental conditions [9, 10, 11]. Although these methods have substantially expanded the circadian analysis toolbox, assessing rhythmicity separately within each experimental group does not directly quantify the magnitude of differences between groups. In particular, inferring differential rhythmicity from the presence or absence of statistically significant rhythms in each group, often summarized using Venn diagrams, is highly susceptible to misclassification and can substantially overestimate physiological effects [10]. Therefore, circadian studies must rely on direct statistical comparisons of rhythmic parameters between groups.

We introduce Circadian Oscillation Detection Analysis and Comparison (CODAC), a computational framework designed to complement existing circadian methods by providing a customizable workflow for rhythmicity estimation and comparison. CODAC comprises four modules: (i) CODAC_single estimates rhythmicity, reports rhythmic parameters, and assigns confidence classifications; (ii) CODAC_flex extends this analysis beyond standard sinusoidal waveforms to detect non-standard rhythmic patterns, including damped and linear-like oscillations; (iii) CODAC_compare enables pairwise comparisons of rhythmicity across two or more groups; and (iv) CODAC_multi accounts for complex experimental designs by clustering multiple groups into different rhythmicity models and assigning a confidence value. Together, CODAC offers a flexible and customizable framework for detecting and comparing rhythmicity across diverse experimental designs. CODAC is freely available as an open-source package at https://github.com/ThiagoPSilveira/CODAC.

## 2 CODAC’s Methodology

The CODAC framework comprises four modules that share a single estimation engine. CODAC_single implements this engine in full: the harmonic model together with its constrained and linearized fits (Section 2.1), the handling of missing and irregularly sampled observations (Section 2.2), the multicriteria rhythmicity score and its four constituent tests (Section 2.3), and the data-driven threshold recommendation (Section 2.4). The remaining three modules are extensions of this common core along two independent axes. CODAC_flex (Section 2.5) generalizes the single characterization fit to a family of candidate waveforms while retaining every inferential and scoring component of CODAC_single. Along the between-group axis, CODAC_compare (Section 2.6) and CODAC_multi (Section 2.7) reuse the CODAC_single engine to fit each experimental group and then, respectively, test for differences in rhythmic parameters between two or more groups and assign a model to the shared-parameter clusters.

### 2.1 Mathematical formulation and parameter estimation

To identify and characterize circadian patterns, CODAC models target expression as a nonlinear harmonic function. The foundational rhythmicity is defined by the standard cosinor equation:

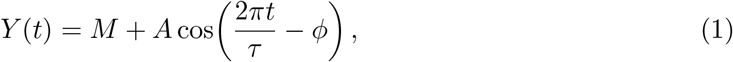

where *Y* (*t*) is the expected expression at time *t*, and the parameters to be estimated are the MESOR *M*, amplitude *A*, period *τ*, and acrophase *ϕ*.

CODAC estimates these parameters through two complementary fits whose outputs serve distinct purposes: a nonlinear constrained fit, which characterizes the oscillation and provides the reported amplitude, acrophase, period, and goodness-of-fit; and a linearized fit, which provides a stable basis for the nested significance test described in Section 2.3.

Parameter estimation is structured as a nonlinear least squares problem. Let *y*_*i*_ be the observed expression at time *t*_*i*_, *i* = 1, …, *n*. The algorithm seeks the parameter vector *θ* = [*M, A, ϕ, τ*]^*T*^ minimizing the Residual Sum of Squares:

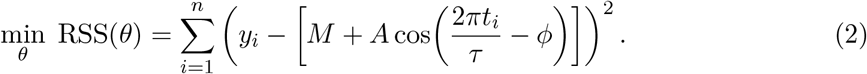

In free-period mode the parameter space is explicitly bounded to exclude physiologically meaningless solutions: the amplitude is non-negative (*A* ≥ 0), the MESOR lies within the empirical range of the data (min_*i*_ *y*_*i*_ ≤ *M* ≤ max_*i*_ *y*_*i*_), the acrophase is constrained to −2*π* ≤*ϕ* ≤ 2*π*, and the period is confined to a physiologically plausible window (default 20 ≤ *τ* ≤ 28 hours). This constrained problem is solved with the Trust Region Reflective (TRF) algorithm. CODAC additionally supports a fixed-period mode, the default, in which *τ* = 24.0 hours is held constant and only *θ* = [*M, A, ϕ*]^*T*^ is estimated by nonlinear least squares; in this mode the non-negativity of the reported amplitude is guaranteed not by an explicit bound but by the curve-based extraction described below, which reads the amplitude as a peak-to-MESOR distance.

To eliminate the “singular gradient” failures common in conventional fitting software, CODAC bypasses arbitrary starting points through a Data-Driven Initialization Heuristic. The initial vector *θ*_0_ is derived directly from the empirical geometry of each target: the MESOR *M*_0_ is the arithmetic mean of the observations, the amplitude *A*_0_ is half the empirical range, the acrophase *ϕ*_0_ is taken from the timepoint of maximum expression, and the period *τ*_0_ is anchored at 24.0 hours. When the period is estimated (free-period mode), the residual surface of (2) is non-convex in *τ* and may contain local minima in which the amplitude collapses toward zero. To guard against such degenerate solutions, CODAC does not depend on a single starting period: it performs a multi-start optimization, re-running the constrained solver from five period seeds spread uniformly across the admissible window and recomputing the acrophase guess *ϕ*_0_ for each seed, and then retains the fit with the highest coefficient of determination. A degenerate local optimum is thus selected only if every seed converges to it, which is exceedingly unlikely in practice. In fixed-period mode there is no period axis to search, so a single fit suffices and the multi-start is not required.

The acrophase that CODAC reports is read from the fitted curve rather than from the raw coefficients: the function is densely evaluated across the sampling window and the acrophase is reported as the time of the resulting peak, which makes it invariant to sign conventions in the parameterization (a fit returning a negative amplitude together with a *π*-shifted acrophase yields the same peak). The amplitude, in contrast, is reported in a model-dependent manner. For the standard cosinor of (1)—the model underlying CODAC_single, CODAC_compare and CODAC_multi—the reported amplitude is the peak-to-MESOR distance of the fitted curve, which for a pure sinusoid equals |*A*| and, read from the curve, is robust to the sign of the fitted coefficient. CODAC_flex, however, additionally fits waveforms carrying a linear baseline trend or an amplitude damping, for which the peak no longer lies a fixed distance *A* above the MESOR; a peak-based measure would absorb the baseline trend in the linear model and overstate the amplitude of the damped models, whose first cycle is the tallest. CODAC_flex therefore reports the amplitude as the fitted amplitude parameter *A* of the winning model itself, the constant amplitude for the standard model, the trend-free oscillation amplitude for the linear model, and the initial amplitude for the damped models, whose subsequent decay is quantified separately by a half-life (see Section 2.5). In every case the amplitude is constrained to be non-negative.

For significance testing, CODAC fits the equivalent linear reparameterization of the cosinor at a fixed period of 24.0 hours,

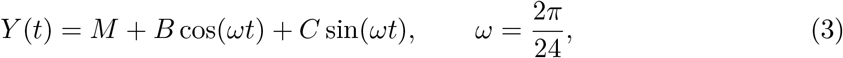

solving for [*M, B, C*]^*T*^ in closed form by ordinary least squares. The amplitude implied by this fit is 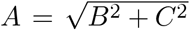; the reported acrophase is not taken from these coefficients but read from the peak of the fitted curve, as described above, so that it is invariant to the sign conventions of the parameterization. Expressing the model linearly yields a numerically stable, three-parameter fit that serves as the alternative hypothesis in the nested *F*-test of Section 2.3.

### 2.2 Handling of missing and unevened sampling time

To handle missing entries (NaNs), CODAC offers three strategies. By default, the engine applies dynamic boolean masking: missing values are identified, and both the TRF solver and the statistical tests are executed strictly on the subset of available points, with the degrees of freedom adjusted on a per-target basis. Targets retaining fewer than four distinct valid timepoints are not fitted and are assigned a non-significant result, preventing overfitting to sparse data. Alternatively, for workflows that require a complete matrix, CODAC optionally offers two other strategies: per-timepoint mean imputation of missing replicates, or strict removal of any target containing missing values. When imputation is selected and a timepoint retains at least one replicate, the missing entries are filled with the mean of the surviving replicates; a timepoint with no surviving replicate is left masked rather than filled, so that the algorithm never invents a value where none was observed. Therefore, masking—proceeding with the available data without imputation or removal—remains the default and recommended behavior.

Because CODAC models observations directly in continuous time, evenly spaced sampling is not required. The framework naturally accommodates irregular sampling intervals, unequal timepoint densities, and asymmetric observation windows.

### 2.3 Multicriteria scoring and rhythmicity classification

CODAC replaces binary rhythm classification with a multicriteria scoring system. Each target is evaluated against four distinct criteria, and one point is awarded for each criterion met:

1. **Statistical significance (***p***-value / FDR)**. The significance of the rhythm is evaluated with a nested *F*-test that compares the linearized cosinor of (3) (three parameters: *M, B, C*) against the null hypothesis of arrhythmicity (a flat line equal to the mean of the observations, one parameter). With *p* = 3 fitted parameters and *n* valid observations, the statistic is

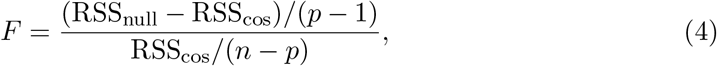

where RSS is the residual sum of squares, *p* − 1 = 2 and *n* − *p* = *n* −3 degrees of freedom, and the *p*-value obtained from the *F* survival function. The user selects the significance threshold (e.g. *α* ≤ 0.05) and the multiple-testing strategy, opting for either raw *p*-values (for hypothesis generation) or the Benjamini–Hochberg (BH) False Discovery Rate (FDR). A point is awarded when the selected *p*-value satisfies the threshold.
2. **Goodness of fit (***R*^2^). The proportion of variance explained by the nonlinear harmonic fit,

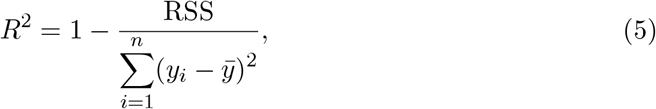

must meet a baseline (default *R*^2^ ≥ 0.4) so that the fitted curve tracks the biological data. The threshold is user-defined. In addition, CODAC computes a data-driven *R*^2^ recommendation via a Support Vector Machine (Section 2.4). This recommendation is for guidance only: it is shown on the diagnostic plots for the user to consider, but the actual scoring uses the threshold set by the user.
3. **Target-specific dynamic amplitude thresholding**. CODAC imposes a dynamically computed minimum amplitude tailored to each target,

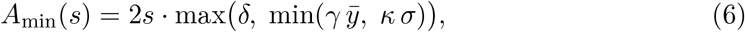

where 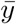 and *σ* are the mean and standard deviation of the per-timepoint means, and the fixed coefficients (*δ* = 0.15, *γ* = 0.10, *κ* = 0.50) impose an absolute technical-noise floor while ensuring a biologically relevant peak-to-trough fold change. The strictness of this criterion is exposed to the user through a single stringency parameter *s* ∈ [0, 1]: the default *s* = 0.5 recovers the validated threshold, *s* = 0 disables the amplitude criterion entirely, and *s* = 1 requires twice the default amplitude. A point is awarded when the fitted amplitude (peak of the curve relative to the MESOR) satisfies *A* ≥ *A*_min_(*s*), and the per-transcript value of *A*_min_ is reported alongside the fitted amplitude so that the comparison is transparent.
4. **Oscillation range**. To distinguish true oscillators from a flat line, CODAC compares the geometric bounds of the fitted curve against an inter-percentile noise band derived from the observed data, after removing per-timepoint outliers via the 1.5 × IQR rule. A Variance Flexibility parameter *ν* ∈ {1, 2, 3} scales the band: the lower and upper percentiles are *P*_low_ = 25 − 10(*ν* − 1) and *P*_high_ = 100 − *P*_low_, yielding the interquartile range (25, 75) at Level 1, (15, 85) at Level 2, and the extreme-prominence band (5, 95) at Level 3. A point is awarded when the fitted curve sweeps beyond this band (its trough falls below *P*_low_ or its peak rises above *P*_high_), empirically demonstrating oscillatory prominence over background noise.

Based on the cumulative score (0–4), each target is mapped to a rhythmicity tier: Arrhythmic (0), Low (1), Medium (2), High (3), or Extremely High (4).

### 2.4 Adaptive threshold recommendation via a support vector machine

Because background stochastic noise varies across experimental designs, a single static *R*^2^ cutoff is difficult to select *a priori*. To assist this choice, CODAC computes a data-driven *R*^2^ recommendation with a linear Support Vector Classifier (SVC).

Within each run, every target with a valid *R*^2^ and *p*-value becomes a training instance. The single feature is the goodness-of-fit *R*^2^, and the binary label is its statistical significance, assigned by comparing the selected *p*-value against the user threshold (*z*_*j*_ = 1 if significant, *z*_*j*_ = −1 otherwise). The linear SVM solves the soft-margin problem

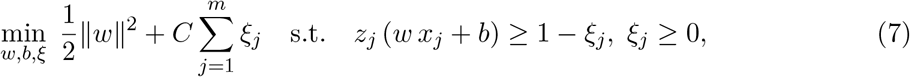

where *x*_*j*_ is the *R*^2^ of the target *j, ξ*_*j*_ are slack variables, and *C* is the regularization parameter. Training proceeds only when both significance classes are present; otherwise the recommendation is skipped.

Once the decision boundary is established, CODAC evaluates the classifier over a dense grid of *R*^2^ values in [0, 1] and reports the smallest *R*^2^ classified as significant as the recommended threshold. In practice, this boundary tends to be lower in high-noise scenarios, detecting valid low-fit oscillators, and higher in low-noise situations, maintaining conservative precision. This value is presented to the user and drawn as a reference line on the *R*^2^ diagnostic plots, complementing rather than replacing the user-defined threshold used in scoring.

### 2.5 CODAC_flex: Extension to non-standard waveforms

CODAC_flex targets datasets whose temporal profiles deviate from a pure sinusoid. It operates on a single experimental group and uses the entire inferential and scoring apparatus of CODAC_single: the nested *F*-test of (4), the four-criterion score and tier mapping of Section 2.3, the adaptive amplitude threshold of (6), the variance-exceedance band, the SVM recommendation of Section 2.4, and the missing-data handling of Section 2.2. Two elements change: the characterization fit of Section 2.1 is replaced by a model competition over a group of candidate waveforms, and the reported amplitude is the fitted amplitude parameter of the winning model, as described in Section 2.1.

#### Candidate waveform family

CODAC_flex fits four models, each reducing to a cosinor oscillation on a possibly non-constant baseline or under a decaying amplitude:

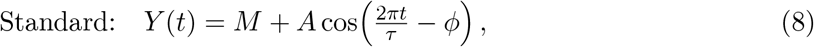

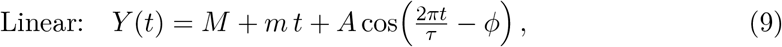

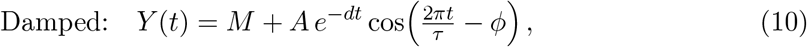

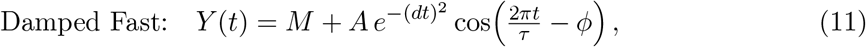

where *m* is a baseline slope and *d* ≥ 0 is a damping rate. The Standard model is (1). The Linear model adds a monotone trend to the MESOR; the Damped and Damped Fast models multiply the oscillation by an exponential (*e*^−*dt*^) and a Gaussian 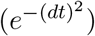 amplitude-decay function, respectively, the latter flattening faster in the tail as expected under de-synchronization-driven amplitude loss. Each model is fitted by constrained least squares (TRF) with the shared CODAC_single bounds (*A* ≥ 0, *M* ∈ [min_*i*_ *y*_*i*_, max_*i*_ *y*_*i*_], −2*π* ≤ *ϕ* ≤ 2*π*, and *τ* [20, 28] hours in free-period mode); the additional parameters are bounded to *m* ∈ [−10, 10] and *d* ∈ [0, 1]. In free-period mode CODAC_flex applies the same five-seed multi-start over *τ* as CODAC_single, independently for each candidate model, keeping each model’s lowest-residual fit before the models compete.

#### Model selection by corrected AIC

The four fitted models are compared by the corrected Akaike Information Criterion. For a model with *k* free parameters fitted to *n* valid observations with residual sum of squares RSS,

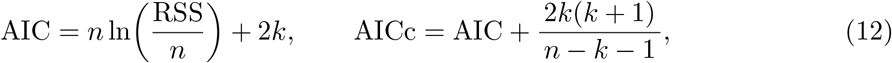

where *k* = 3 for the Standard model and *k* = 4 for the Linear, Damped, and Damped Fast models (each increased by one when the period is free); the finite-sample correction term reverts to plain AIC when *n* − *k* − 1 ≤ 0. The winning model is the one with the lowest AICc, so that the added flexibility of a trend or a damping parameter is retained only when it improves the fit enough to offset its extra parameter.

#### Reported amplitude and damping

The reported amplitude is the fitted parameter *A* of the winning model (Section 2.1): the constant amplitude for the Standard model, the trend-free oscillation amplitude for the Linear model, and the initial amplitude for the damped models. For the damped models the amplitude decay is summarized by the damping half-life,

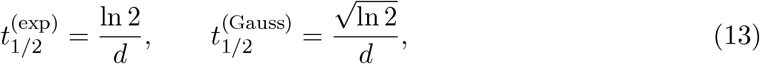

for the Damped and Damped Fast models, respectively. All remaining outputs (rhythmic parameters, *R*^2^, *p*-value/FDR, and the multicriteria tier) are produced exactly as in CODAC_single.

### 2.6 CODAC_compare: Differential rhythmicity between two groups

CODAC_compare addresses two experimental groups, or pairwise comparisons among more than two groups, and asks not whether each is rhythmic in isolation but whether their rhythmic parameters differ. It inherits, without modification, the per-group machinery of CODAC_single and adds a differential layer.

#### Per-group characterization (inherited)

Each of the two groups is characterized independently with the standard cosinor engine of Sections 2.1 to 2.4: the same constrained and linearized fits, nested *F*-test, *R*^2^, adaptive amplitude threshold, variance-exceedance band, SVM recommendation, and missing-data handling. This yields, for each group, its MESOR, amplitude, acrophase, and period together with its multicriteria rhythmicity tier. CODAC_compare uses only the standard cosinor model of (1).

#### Rhythmicity structure

A target’s rhythmicity structure is fixed by the per-group tiers, not by the difference tests. Under a user-selected rhythmicity cutoff (default High), each group is deemed rhythmic or not, and the pair is labelled *both rhythmic, group 1 only, group 2 only*, or *neither rhythmic*. This structural marker is computed once and is independent of the pairwise *p*-values, so the significance-driven sub-decisions below can be recomputed from raw or FDR *p*-values without refitting.

#### Differential parameter tests

The magnitude and significance of each parameter difference are assessed by a nested nonlinear test on the pooled observations of the two groups. Let *g* ∈ {0, 1} indicate group membership (0 reference, 1 comparison) and fix the angular frequency at *ω* = 2*π/τ*_ref_, with *τ*_ref_ the mean of the two per-group period estimates. The full model lets each rhythmic parameter shift between groups,

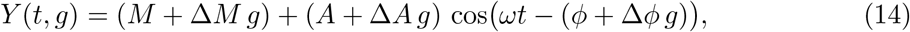

and three restricted models each null one differential parameter (Δ*M* = 0, Δ*A* = 0, or Δ*ϕ* = 0). Fitting the full (6-parameter) and each restricted (5-parameter) model by nonlinear least squares, and denoting their residual sums of squares RSS_full_ and RSS_r_, the significance of each difference follows from an extra-sum-of-squares *F*-test with 1 and *n* − 6 degrees of freedom,

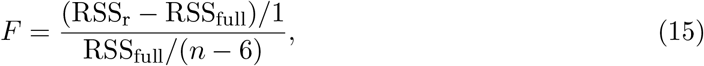

yielding the per-parameter *p*-values *p*_Δ*M*_, *p*_Δ*A*_, and *p*_Δ*ϕ*_ (the test requires a total of *n* > 6 individual data points pooled across both groups). The MESOR difference is always tested. The amplitude difference is evaluated when at least one group exhibits rhythmicity. In pairs where both groups are rhythmic, it measures the amplitude change; in pairs with only one rhythmic group, it assesses the confidence in the apparent rhythm loss or gain. Meanwhile, phase difference is tested only when both groups are rhythmic, as phase cannot be meaningfully interpreted if one group is arrhythmic. Reported alongside are the raw parameter differences Δ*M* = *M*_1_ − *M*_2_, Δ*A* = *A*_1_ − *A*_2_, and the circular phase difference wrapped to [−*τ*_ref_/2, *τ*_ref_/2].

#### Omnibus tests and classification

CODAC_compare first reports two omnibus extra-sum-of-squares *F*-tests from the linearized cosinor with group interactions: overall rhythmicity (any group rhythmic versus none) and overall rhythm difference. CODAC_multi additionally uses an orthogonal pooled shared-rhythm test to screen the multiple-testing correction (see Section 2.7). Parameter differences are then classified at significance level *α*. On the *MESOR* axis, *p*_Δ*M*_ defines *Different, Conserved*, or *Undetermined*. On the *rhythm* axis, neither-rhythmic pairs are Category 1, singly-rhythmic pairs are Category 2 or 3 and receive a loss/gain confidence flag based on *p*_Δ*A*_, and both-rhythmic pairs are Category 4–7 according to the significance of *p*_Δ*A*_ and *p*_Δ*ϕ*_: unchanged, amplitude change, phase change, or both, respectively.

#### Multiple-testing correction

Within each pair, the three per-parameter *p*-values are corrected for multiple testing by the BH procedure, each component forming its own family across targets. Furthermore, all global omnibus tests (such as the global rhythmicity test and global difference test) are also independently corrected across all targets.

### 2.7 CODAC_multi: Rhythmicity grouping across multiple groups

Following the analytical frameworks of the previous modules, CODAC_multi handles more complex experimental conditions by generalizing the two-group comparison to an arbitrary number of *k* groups. Rather than only testing differences pair by pair, CODAC_multi evaluates how the groups cluster into sets that share a common rhythm or a common MESOR, selecting the optimal clustering through information-criterion model selection.

#### Two independent clustering tiers

Beyond the pairwise analysis, CODAC_multi performs separate clustering on the rhythm and baseline (MESOR) axes, treating changes in oscillation and overall level as distinct effects. Each clustering is selected by model comparison across group partitions. For each candidate partition, an OLS model is fitted at a fixed 24-hour period, with shared harmonic terms (cos *ωt*, sin *ωt*) within rhythm blocks or a shared intercept within MESOR blocks, while the other axis remains free by group as a nuisance component.

#### Information criterion

With residual sum of squares RSS, *n* observations, and *k* mean-structure design columns, the criterion is, by default, the Schwarz information criterion

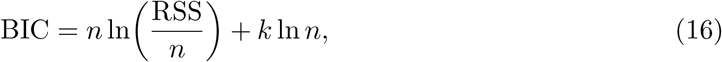

with an optional corrected-AIC alternative AICc = *n* ln(RSS*/n*) + 2*k* + 2*k*(*k* + 1)/(*n* −*k* −1) sharing the convention of (12) (the variance is not counted as a parameter). BIC is the default because its heavier complexity penalty is more conservative about declaring a difference and curbs the over-splitting to which AICc is prone when a rhythm or a baseline is in fact shared. The selected partition minimizes the criterion.

#### Confidence

The relative support for the winning partition is reported as its Schwarz weight,

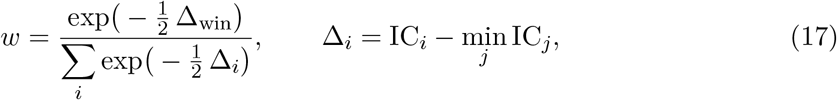

together with the criterion gap between the best and second-best partitions. This weight is defined only when a partition search is actually run (Section 2.7); where the outcome is fixed without a search, no weight is reported.

#### Four omnibus tests

The clustering is anchored by a set of nested extra-sum-of-squares *F*-tests built from three ordinary least squares fits at *ω* = 2*π/*24 (the fourth, baseline test uses a separate nested pair): a group-specific-mesors-only model ℳ_0_; a shared-rhythm model ℳ_1_ that adds one pair of harmonic terms (cos *ωt*, sin *ωt*) common to all groups; and a pergroup-rhythm model ℳ_2_ that lets the harmonic terms vary by group. Comparing them yields

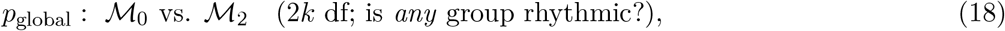

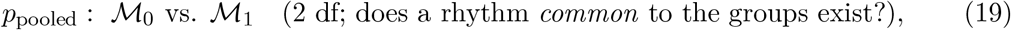

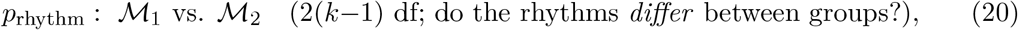

so that the 2*k* degrees of freedom (df) of the global test decompose into the 2 of the pooled shared-rhythm effect and the 2(*k* −1) of the group-by-rhythm interaction; for balanced sampling these two components are orthogonal, and the permutation calibration (`permute_B`) confirms that the gate’s empirical false discovery rate is controlled even when sampling is unbalanced. A fourth test, *p*_mesor_, is an omnibus baseline test, a nested *F*-test of an all-shared-baseline model against an all-free-baseline model, controlling for a free per-group rhythm. Each of the four is also reported BH corrected across all targets in a companion column; both the raw and corrected values are always retained.

#### Gating of the two tiers

On the rhythm axis, the rhythmic set *R* contains the groups that meet the rhythmicity cutoff (default High) according to the per-group multicriteria tier of Section 2.3. The overall test *p*_global_ is reported for context but does not determine group rhythmicity. The partition search is restricted to *R* and is performed only when the rhythm-difference gate is significant: for |*R*| ≥ 2, partitions are searched when *p*_rhythm_ ≤ *α*, whereas *p*_rhythm_ > *α* yields a single shared-rhythm block. No search is required for |*R*| ≤ 1, and an untestable *p*_rhythm_ leaves the grouping *undetermined*. Similarly, the MESOR partition is searched only when *p*_mesor_ ≤ *α*; otherwise, a common baseline is assigned to all groups. The corrected *p*_rhythm_ used at the gate is selected by the screening procedure described next.

#### Power-preserving screened correction

Correcting *p*_rhythm_ across all targets can be overly conservative in circadian screens, where most targets are often arrhythmic. CODAC_multi therefore uses a two-stage correction that exploits the independence established in (19)–(20). First, *p*_pooled_ is BH-corrected across all targets, and only targets passing the *α* threshold are retained. Within this screened set, *p*_rhythm_ is BH-corrected again. Because the pooled-rhythm test is independent of the group-by-rhythm interaction, this screening does not bias the interaction test and typically provides substantially greater power. The main blind spot is near-antiphase groups, whose rhythms may cancel in the pooled test. For this reason, the genome-wide correction is always reported alongside the screened correction. A switch determines which corrected value controls the gate, with the genome-wide correction as the conservative default.

#### Empirical calibration by permutation

Because the screened correction is a two-stage procedure, CODAC_multi provides an optional permutation diagnostic that measures its realized error rate in the engine itself. Group labels are permuted within each timepoint, a null in which the groups share the common rhythm but carry no systematic between-group difference, and the four omnibus tests, the screen, and the correction are recomputed on each permuted replicate. Counting how many targets the gate declares different under the null, relative to the number declared on the observed data, gives an empirical false discovery rate for the gate.

#### Reconciliation of the pairwise category

In CODAC_compare, the seven-category rhythm label of Section 2.6 is assigned independently to each pair. In CODAC_multi, the omnibus clustering is authoritative, preventing conflicts between pairwise and multi-group classifications. For pairs in which both groups are rhythmic, groups in the same rhythm block are assigned Category 4 (unchanged), whereas groups in different blocks are assigned Category 5, 6, or 7 according to the pairwise amplitude and phase tests (*p*_Δ*A*_, *p*_Δ*ϕ*_) in (15). Thus, shared blocks yield Category 4 within blocks and Category 5–7 between blocks. Pairs involving an arrhythmic group retain their loss/gain category from Section 2.6. This reconciliation may override an isolated pairwise significance result, but all raw differences and their *p*-values remain in the output.

#### Model space, codes, and scaling

A rhythm model assigns the groups to rhythmic blocks while allowing some groups to remain arrhythmic; a MESOR model partitions all groups into baseline blocks, since every group has a baseline. Rhythm models are labelled M01, M02, … and MESOR models MM1, MM2, …, following a fixed order based on the number and identity of rhythmic groups and, finally, from the coarsest to the finest partition. These codes are stable for a fixed number of groups and can therefore be compared across targets and runs of the same design, but not across designs with different *k*; grouping structure, by contrast, is design-independent. The model space grows exponentially, with Bell(*k*+1) rhythm models and Bell(*k*) MESOR models.

## 3 CODAC’s benchmarking

### 3.1 Automated topographical visualization

CODAC automatically generates graphical summaries to facilitate the interpretation of rhythmicity results. Phase distributions are visualized using rose plots. Heatmaps also display rhythmic targets ordered by acrophase, with expression values standardized across time to emphasize differences in amplitude and phase. For CODAC_compare, heatmaps are generated according to the assigned rhythmicity categories (1 to 7) and display the temporal profiles of the compared groups. CODAC_multi additionally generates model-specific and consolidated heatmaps, allowing groups with shared or distinct rhythmic patterns to be visualized according to their selected grouping model.

### 3.2 *In silico* circadian dataset benchmark

CircaInSilico software [12] was used to generate *in silico* circadian datasets with varied numbers of replicates and sampling intervals. Additional settings included a fixed period of 24 hours, minimum and maximum amplitudes of 1 and 6, respectively, an outlier amplitude set to zero, 2,000 observations per dataset, and a rhythmicity-positive rate of 50%. The generated datasets were used to benchmark the CODAC_single and CODAC_flex functions, employing the standard model and an interval variance flexibility of 1. The minimum *R*^2^ threshold was set to 0.4 when the SVM recommendation was not used or adjusted to the *R*^2^ values suggested by the SVM. Missing values were handled using the `KEEP` function. For the bioluminescence-data benchmark, simulated data were generated using the following equation:

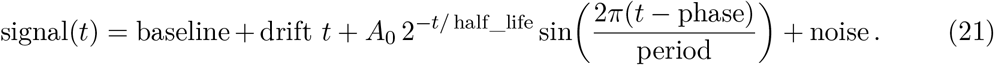

The simulated dataset contained 10 biological replicates sampled at 1-hour intervals for 120 hours. CODAC_flex was run using the same configuration as CODAC_single, but the period was allowed to vary between 20 and 28 hours.

### 3.3 Real-data circadian dataset benchmark for CODAC_single

Rhythmicity was assessed across five computational methods applied to the same normalized expression datasets. All methods assumed a fixed circadian period of 24 hours. BH multiple-testing correction was applied independently within each method.

#### 3.3.1 CircaSingle

The `circa_single` function from the CircaCompare package [9] was used to fit each gene independently using a nonlinear 24-hour cosinor model. All replicate observations across Zeitgeber time (ZT) points were pooled, and fitting was performed without an internal rhythmicity filter by setting `alpha_threshold = 1`. A timeout of 10,000 was set for each gene. The period was fixed at 24 hours. Rhythmicity was assessed using the rhythmicity *p*-value returned by `circa_single`, followed by BH correction.

#### 3.3.2 DryR

The `f24` function from the DryR package was used because the data were already normalized. The input consisted of a gene-by-sample expression matrix with a corresponding time vector containing all replicate ZT observations and a fixed period of 24 hours [11].

#### 3.3.3 Classical cosinor analysis

Classical cosinor analysis was performed using the `cosinor.lm` function from the R package `cosinor` (https://www.rdocumentation.org/packages/cosinor/versions/1.2.3). Each gene was fitted independently using all replicate observations and a fixed period of 24 hours. Rhythmicity was assessed using the BH-corrected amplitude *p*-value, corresponding to the null hypothesis of zero amplitude.

#### 3.3.4 GLMMcosinor

GLMMcosinor [13] was used. Each gene was modeled independently in long format, retaining all replicate observations at each timepoint. For each gene, expression was fitted as a function of time using a 24-hour cosinor model with Gaussian errors through `cglmm()` and `glmmTMB`. Rhythmicity was tested using the amplitude *p*-value, corresponding to the null hypothesis of zero amplitude. BH-corrected amplitude *p*-values were used to estimate rhythmicity.

#### 3.3.5 CODAC_single

CODAC_single was used with the standard parameters: a fixed period of 24 hours, a minimum *R*^2^ of 0.4, a *p*-value threshold of 0.05, a minimum rhythmicity-probability category of `HIGH`, and an interval variance flexibility of 1. Missing values were handled using the `KEEP` function.

#### 3.3.6 Comparison across algorithms

Across all algorithms, rhythmic-gene sets were compared at BH-adjusted *p*-value thresholds of 0.01, 0.05, 0.10, and 0.20.

### 3.4 Reanalysis of public circadian datasets for CODAC_single

Public transcriptome data were obtained from previous studies using different sampling intervals. For the dataset sampled every 6 hours, data were extracted and normalized as described in the original publication [14]. The same workflow was applied to the dataset sampled every 4 hours, although the RNA-seq method used in that study was 3^*′*^ RNA-seq [15]. For the microarray dataset [16], probes targeting the same gene were averaged and the data were log_2_-transformed. Lowly expressed genes were defined as those below the 25th percentile, resulting in 15,975 genes.

### 3.5 Reanalysis of public circadian datasets for CODAC_compare

Public transcriptome data were obtained from several publications [15, 17, 18]. For each dataset, genes with a total read count ≤ 100 across all samples were removed, except for the dataset from Deota et al., for which a threshold of ≤ 10 was used [18]. A common experimental design specifying condition, sampling time, and a 24-hour period was used for all analyses.

DryR and CompareRhythms were run on raw counts. DryR applied its model-selection framework, with classifications accepted when the Bayesian information criterion weight (BICW) was greater than 0.4. CompareRhythms was run using the DESeq2 option, with rhythmicity and comparison FDR thresholds of 0.05 and an amplitude cutoff of 0.5. For CircaCompare, diffCircadian, and CODAC, counts were transformed using the DESeq2 variance-stabilizing transformation. CircaCompare was fitted separately to each gene to estimate group-specific rhythmicity and differences in amplitude and phase. diffCircadian first assessed rhythmicity in each condition using `LR_rhythmicity`, followed by `LR_diff` tests for amplitude in genes rhythmic in either condition and for phase in genes rhythmic in both conditions. CODAC was run using an *R*^2^ threshold of 0.4, a significance threshold of 0.05, high-confidence rhythmicity criteria, and an amplitude-stringency parameter of 0.5. Analyses including and excluding medium-probability rhythmic targets were evaluated.

Because each method reports different native classifications, the results were translated into a common vocabulary: Arrhythmic, Gain, Loss, Same, differentially rhythmic gene (DRG), and Ambiguous. Gain and Loss indicated rhythmicity restricted to the reference condition and required supporting evidence of an amplitude difference or a high-confidence method-specific classification. Genes rhythmic in only one condition without an amplitude change were classified as Ambiguous. Genes rhythmic in both conditions were classified as DRGs when amplitude and/or phase differed significantly. Changes in MESOR were not used to define the harmonized categories. Low-confidence DryR classifications (BICW < 0.4) were assigned as Ambiguous, whereas genes omitted by the CompareRhythms rhythmicity screen were considered Arrhythmic.

Pairwise agreement between methods was assessed using Cohen’s *κ* applied to the harmonized categorical classifications. For each pair of methods, the analysis included only genes with non-missing calls from both methods, and the number of genes contributing to each comparison was recorded. Cohen’s *κ* accounts for agreement expected by chance based on the marginal frequency of each classification category.

## 4 Results

### 4.1 CODAC_single *in silico* benchmarking

The initial validation of CODAC_single was conducted using simulated datasets with the period fixed at 24 hours and the proportion of rhythmic observations set to 50%. At a sampling interval of 2 hours, CODAC_single effectively detected rhythms using both uncorrected and BH-adjusted *p*-values. Rhythm detection was not strongly affected by increasing the number of biological replicates. Under the 2-hours sampling condition, the Medium category produced an apparent detection rate above 100%. This over-detection decreased as the number of replicates increased. Under these conditions, incorporation of the support vector machine (SVM) recommendation had a negligible effect on the overall rhythmicity-detection rate (Figure 1, left panel).

**Figure 1:**
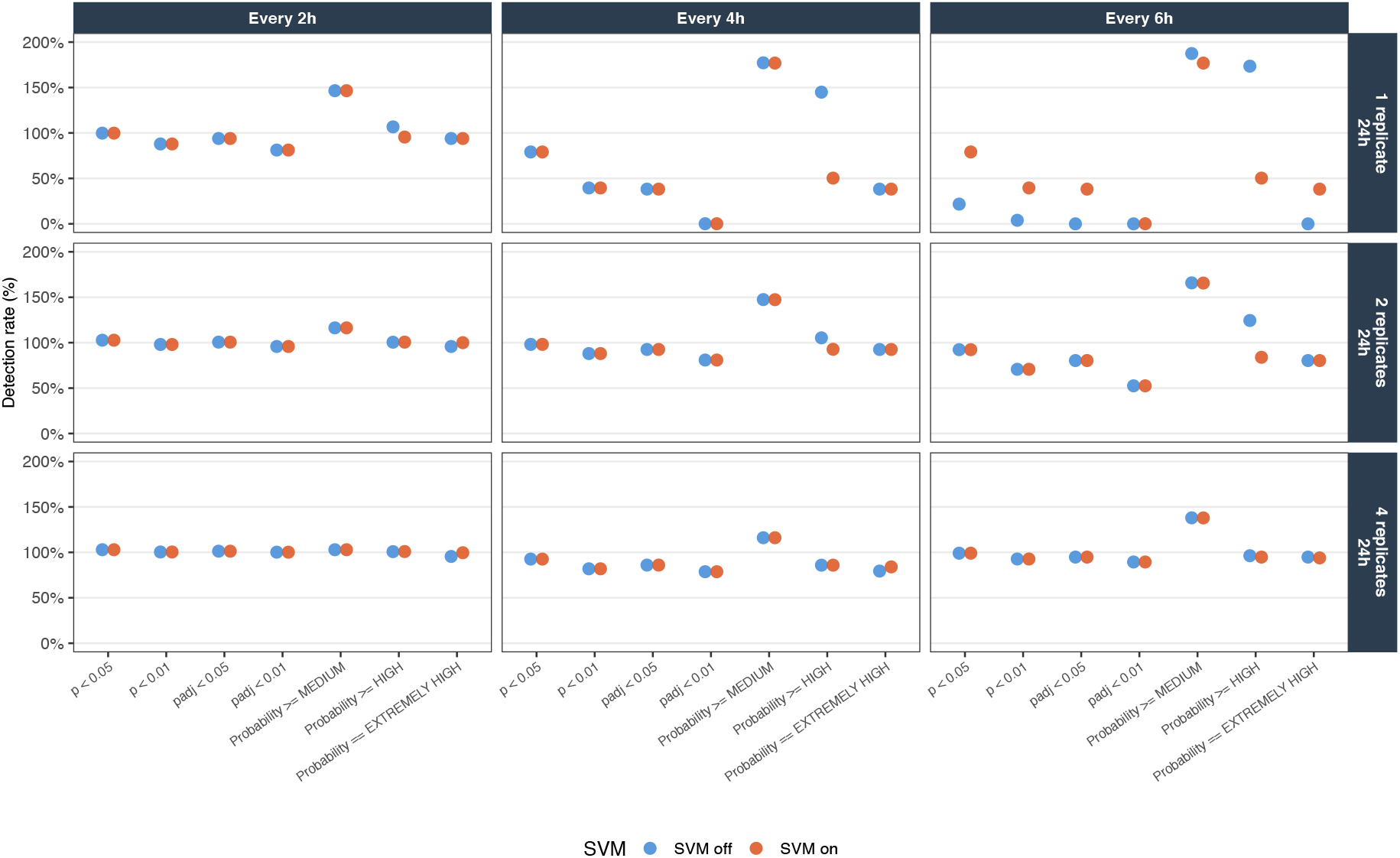
CODAC_single benchmarking across different sampling intervals and replicate numbers. Simulated *in silico* circadian datasets were analyzed using different sampling intervals, replicate numbers, rhythmicity-confidence categories, and the presence or absence of the SVM-derived *R*^2^ recommendation. The left, middle, and right panels show results obtained with sampling intervals of 2, 4, and 6 hours, respectively.

When the sampling interval was increased to 4 hours, the use of a single replicate substantially compromised the ability of CODAC_single to detect rhythms, regardless of the analytical parameters applied. The Medium category again produced the highest detection rate and was generally unaffected by SVM incorporation, except within the High-probability category (Figure 1, middle panel). Increasing the number of replicates per timepoint restored detection rates to approximately 100% under both the two-replicate and four-replicate conditions. The SVM recommendation produced only a marginal effect under these conditions.

Extending the sampling interval to 6 hours while retaining only one replicate markedly reduced the rhythm-detection rate. The SVM recommendation provided some improvement, particularly in the High and Extremely High rhythmicity categories when one or two replicates were available (Figure 1, right panel). At the 6-hours sampling interval, increasing the number of replicates to at least two restored CODAC_single detection rates to values close to 100%. The Medium category again showed the highest apparent detection rate, exceeding 100%, although this effect decreased as the number of replicates increased.

Across all tested conditions, rhythmic genes identified by CODAC_single showed substantial overlap with the rhythmic genes contained in the original simulated datasets. Even under the lowest-resolution sampling condition, with measurements collected every 6 hours, the overlap was at least 85%. These findings indicate that most simulated rhythmic genes were consistently detected by CODAC_single.

### 4.2 CODAC_single real-data benchmarking

The performance of CODAC_single was evaluated using public circadian transcriptomic datasets, including RNA-seq and microarray datasets collected at different sampling intervals (see material and methods section). CODAC_single was compared with established rhythmicity-detection methods, including classical Cosinor, CircaSingle from CircaCompare, `f24` from DryR, GLMMcosinor, and diffCircadian. Because the selected methods are based on sinusoidal model fitting, non-parametric methods designed to identify non-sinusoidal rhythmic patterns, such as JTK_CYCLE and RAIN, were not included in the comparison. Although CircaN [19] also uses sinusoidal fitting, it applies a different fitting strategy that accommodates additional waveform types and was therefore not included in this benchmark.

Real-data benchmarking was performed using datasets with sampling intervals ranging from 2 to 6 hours. The first analysis used a mouse-liver circadian dataset consisting of young male and female mice maintained on a chow diet, with samples collected every 6 hours [14]. CODAC_single performed similarly to DryR and diffCircadian and consistently identified a comparable number of rhythmic genes. When the CODAC multicriteria classification was set to High, relaxing the false discovery rate (FDR) cutoff slightly increased the number of rhythmic genes detected. Incorporation of the SVM-derived *R*^2^ recommendation provided no additional benefit for rhythm detection. In contrast, classical Cosinor, CircaSingle, and GLMMcosinor identified almost twice as many rhythmic genes, even when a stricter threshold of FDR < 0.01 was applied. When the FDR threshold was relaxed, this difference increased, with more than 10,000 genes classified as rhythmic at FDR < 0.20. Each of these three methods also identified method-specific rhythmic-gene sets, contributing to the larger number of rhythmic genes detected at FDR < 0.05 (Figure 2B).

**Figure 2:**
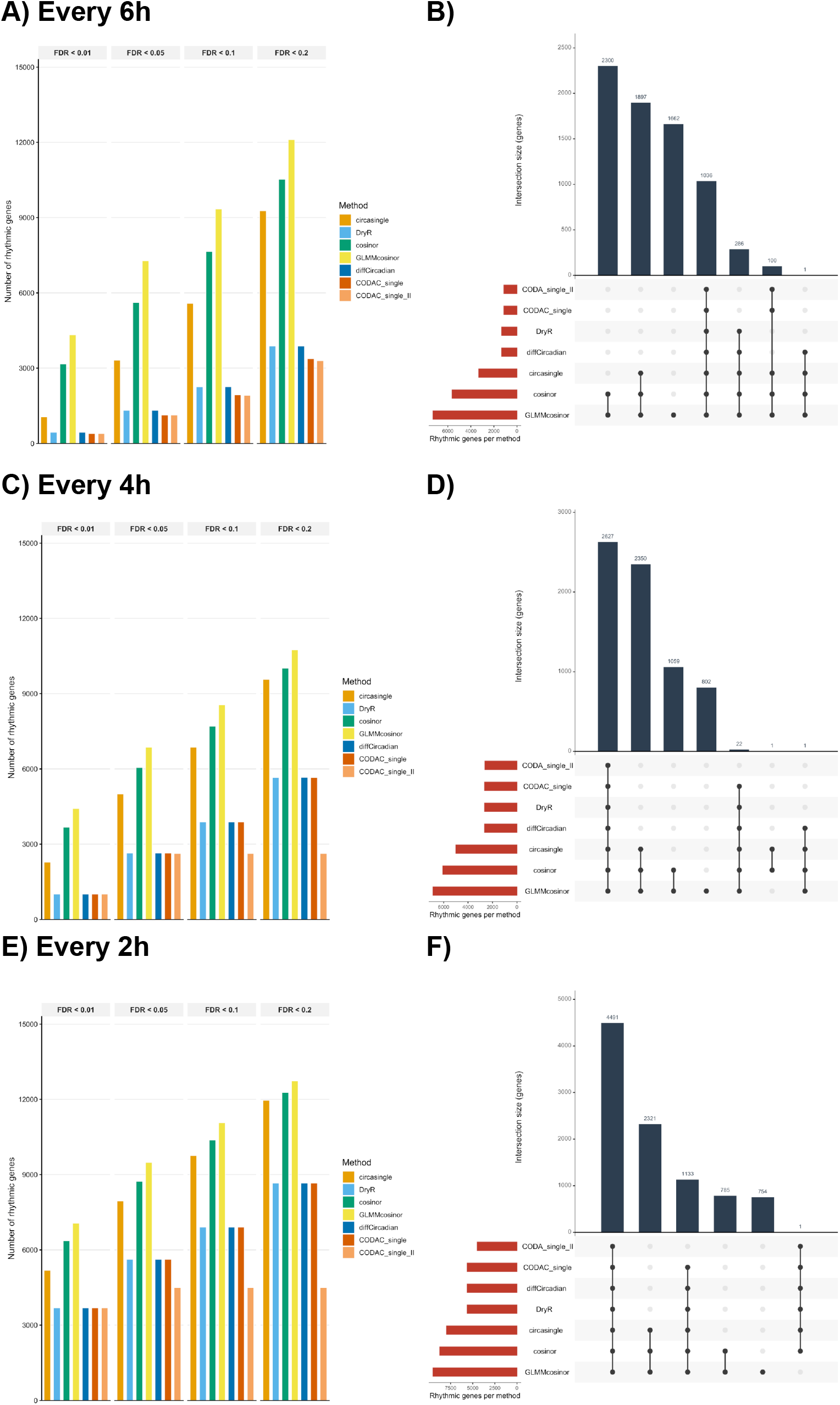
CODAC_single detects rhythmicity in real datasets at rates comparable to established circadian algorithms. **A, C, and E)** Bar plots show the numbers of rhythmic genes identified by the evaluated algorithms at different FDR thresholds in datasets collected using different sampling intervals. **B, D, and F)** UpSet plots show the overlap among the rhythmic-gene sets identified by the evaluated methods. CODAC_single represents genes identified using adjusted *p*-value thresholds alone, whereas CODAC_single_II represents genes satisfying at least the High rhythmicity-confidence classification.

The second benchmark used liver data from young female and male *Bmal1* ^lox/lox^ mice maintained on a chow diet [15]. Under this condition, DryR, CODAC_single, and diffCircadian showed similar performance across the evaluated FDR thresholds. Genes assigned to the High rhythmicity category by CODAC_single were evaluated using a fixed threshold of FDR < 0.05. CircaSingle, classical Cosinor, and GLMMcosinor detected larger numbers of rhythmic genes even under more restrictive FDR thresholds. This difference approximately doubled when the FDR threshold was relaxed beyond 0.10. Across all methods, 2,627 genes were consistently identified as rhythmic (Figure 2C–D). The methods were also evaluated using a landmark mouse-liver dataset sampled every 2 hours over three circadian cycles [16]. The overall performance pattern remained similar, without major differences from those observed in the lower-resolution datasets (Figure 2E–F).

Overall, these findings demonstrate that CODAC_single detects rhythmic genes at rates comparable to established approaches such as DryR and diffCircadian. The High multicriteria rhythmicity classification provided an effective and conservative definition of rhythmicity at an FDR threshold of 0.05.

### 4.3 CODAC_flex

CODAC_flex was developed as an alternative to the standard CODAC_single model for datasets containing more diverse temporal patterns. In addition to the standard sinusoidal model, CODAC_flex supports linear, damped, and rapidly damped waveform models. The rapidly damped model is represented in the software output as `damped_fast`. For each gene or temporal profile, CODAC_flex fits the available candidate models and selects the model providing the best fit. Model-specific parameters are also reported. For damped rhythms, these outputs include the estimated damping rate and half-life, allowing the progressive reduction in oscillation amplitude to be quantified and visualized.

Using the high-resolution mouse-liver dataset sampled every 2 hours [16], CODAC_flex classified approximately 85.9% of genes as best represented by the standard sinusoidal model. The remaining genes were assigned to linear models (9.4%) or damped models (4.7%). Representative examples of each waveform category are shown in Figure 3.

**Figure 3:**
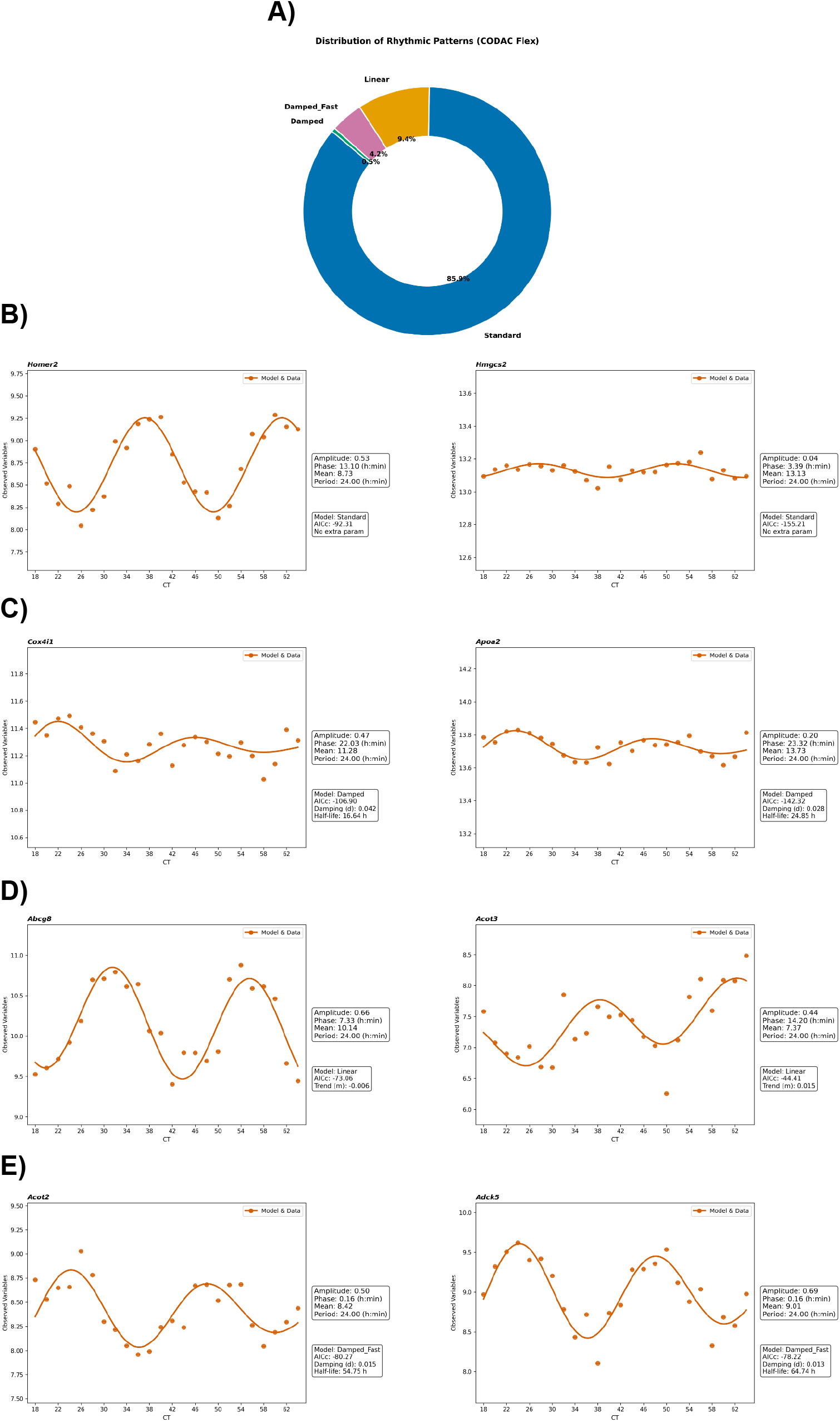
CODAC_flex detects non-standard rhythmic waveforms in a high-resolution microarray dataset. **A)** Distribution of temporal profiles assigned to the candidate waveform models. **B)** Representative profiles classified using the Standard model. **C)** Representative profiles classified using the Damped model. **D)** Representative profiles classified using the Linear model. **E)** Representative profiles classified using the Damped Fast model.

Estimating damping is especially relevant for long-duration *in vitro* and *ex vivo* bioluminescence experiments. These experiments may use clock-protein fusion reporters, such as PER2::LUC, or circadian promoters controlling luciferase expression, such as BMAL1::LUC. In these systems, oscillation amplitude frequently decreases with each successive cycle, making the damping rate an important experimental parameter. Using a synthetic bioluminescence dataset, CODAC_flex successfully identified damped rhythms while allowing the circadian period to vary between 20 and 28 hours (Figure 4).

**Figure 4:**
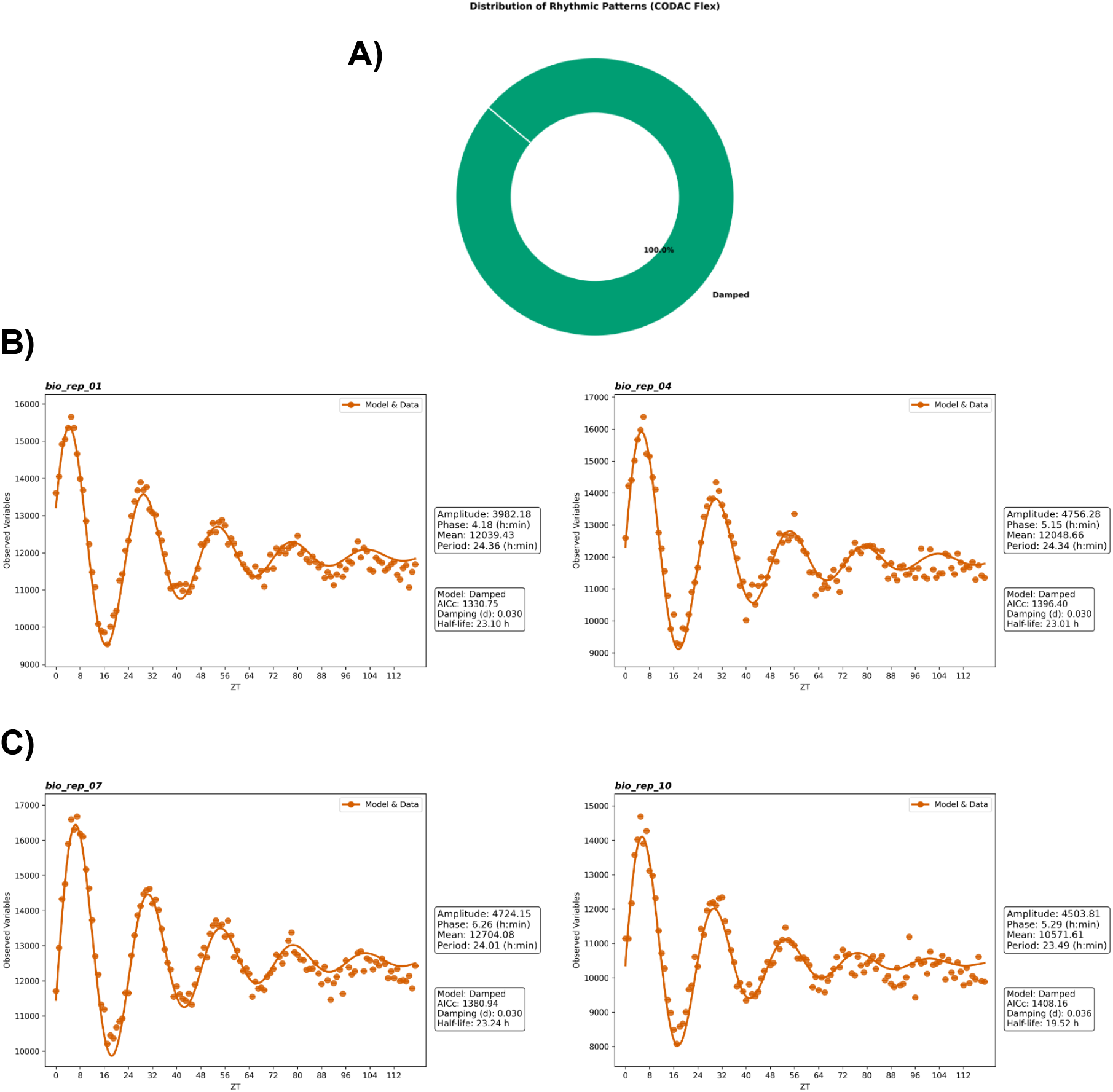
CODAC_flex detects damped rhythms in simulated high-resolution bioluminescence data. **A)** Distribution of the waveform models assigned to the simulated profiles. **B–C)** Representative simulated bioluminescence profiles and their corresponding fitted curves.

Overall, CODAC_flex provides an alternative method for evaluating temporal profiles that deviate from standard sinusoidal oscillations. It is particularly suitable for high-resolution datasets and experimental systems in which progressive damping is expected.

### 4.4 CODAC_compare

CODAC_compare was designed to compare rhythmicity between two experimental groups while distinguishing changes in rhythmic parameters from changes in MESOR. For each comparison, targets are assigned to one of seven biological categories.

Category 1 comprises targets that are arrhythmic in both groups. Categories 2 and 3 identify targets that are rhythmic in only one group and therefore represent potential losses or gains of rhythmicity. These classifications are considered high confidence only when the between-group amplitude difference is significant according to either the uncorrected or BH-adjusted *p*-value. When the amplitude difference is not significant, the classification is interpreted as weak evidence for a gain or loss of rhythmicity. Category 4 includes targets that are rhythmic in both groups without significant differences in amplitude or phase and therefore represents conserved rhythmicity. Categories 5–7 describe differential rhythmicity among targets that are rhythmic in both groups: amplitude changes only, phase changes only, or combined amplitude and phase changes, respectively. The corresponding parameter-difference estimates quantify the magnitude and direction of each change.

Targets assigned to the Medium rhythmicity-confidence category may be retained for exploratory analyses or excluded to reduce uncertainty and the multiple-testing penalization. For confirmatory analyses, we recommend excluding Medium-confidence targets. MESOR is evaluated independently of the seven rhythmicity categories, allowing changes in average expression to be distinguished from changes in amplitude, phase, or rhythmicity status.

To evaluate the performance of CODAC_compare, we reanalyzed 11 publicly available transcriptomic datasets using a uniform analytical pipeline and harmonized the method-specific nomenclature used to describe changes in rhythmicity. Pairwise agreement between CODAC_compare and DryR, CompareRhythms, CircaCompare, and diffCircadian was assessed using Cohen’s *κ*. Agreement was consistently higher for arrhythmic classifications than for rhythmic classifications. For arrhythmic calls, mean *κ* values generally indicated moderate-to-substantial agreement, with the strongest concordance observed between CODAC_compare and diffCircadian (Figure 5A). Agreement for rhythmic classifications was lower and more variable across methods and statistical thresholds, indicating only partial concordance in the identification and classification of rhythmic genes (Figure 5B). DryR generally showed the lowest agreement with CODAC_compare. Excluding medium-confidence rhythms consistently increased Cohen’s *κ* for both arrhythmic and rhythmic classifications.

**Figure 5:**
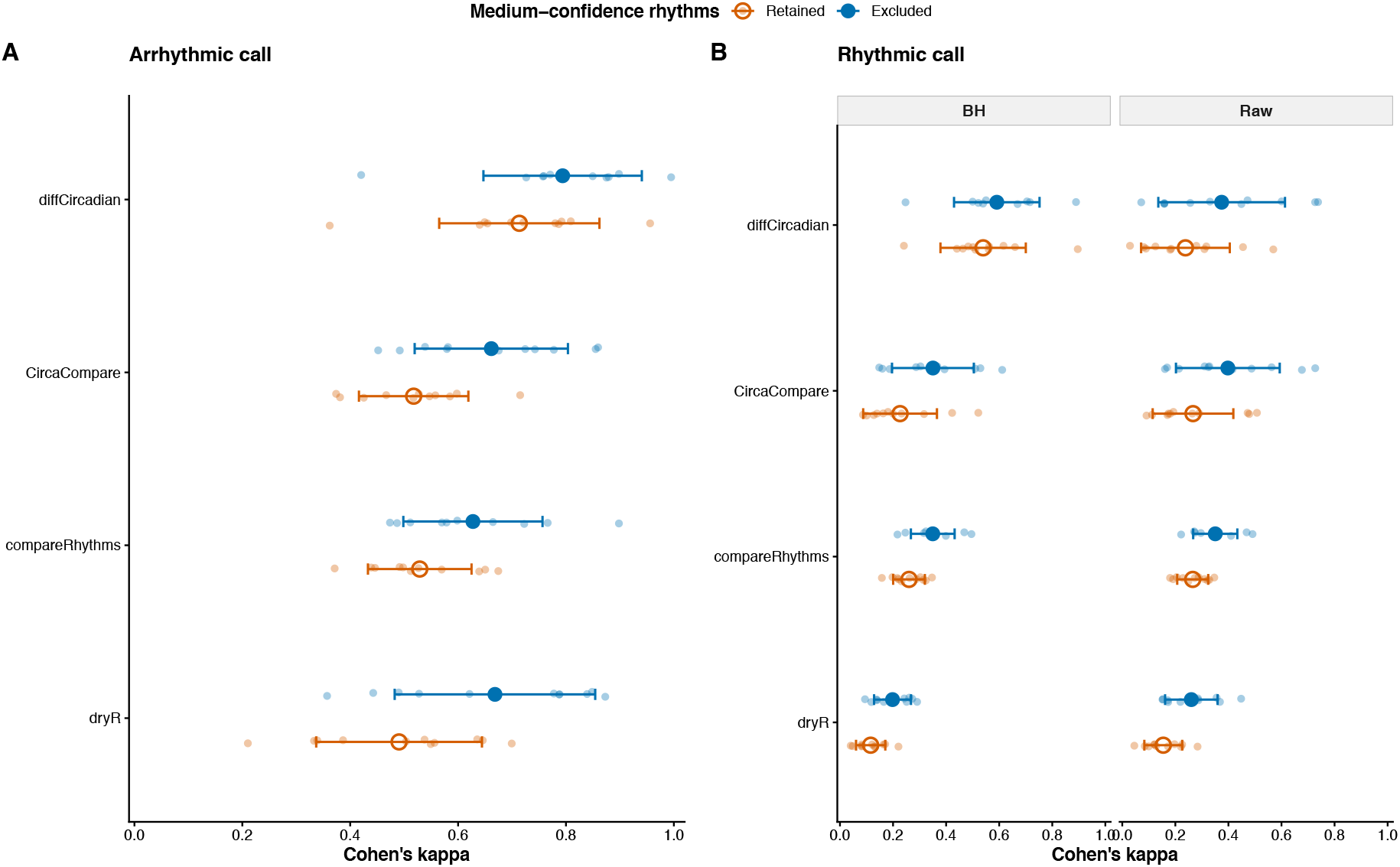
Agreement between CODAC_compare and established differential-rhythmicity methods. Pairwise agreement was evaluated across 11 public transcriptomic datasets using Cohen’s *κ*. **A)** Agreement for arrhythmic classification. **B)** Agreement for rhythmic classifications. Analyses were performed both with and without targets assigned to the Medium rhythmicity-confidence category.

This result demonstrates that borderline rhythmicity calls were an important source of disagreement among methods and supports the exclusion of medium-confidence targets when a more conservative comparison is required.

### 4.5 CODAC_multi

CODAC_multi was developed to compare rhythmicity across several experimental groups and identify the most likely model of change. Rhythmicity and rhythmic parameters are first estimated within each group using the CODAC_single framework. Four global tests are then computed for each target: whether any group is rhythmic, whether a rhythm common to the groups exists, whether the rhythms differ between groups, and whether the baselines differ. All tests are BH-corrected.

On the basis of these tests, CODAC_multi searches the full space of configurations in which the groups can share a rhythm and selects the one with the best information criterion (either BIC or AICc). The result is reported on two independent axes: which groups share the same rhythm, and which share the same baseline (MESOR). A group enters a rhythmic block only if it passes CODAC’s multi-criteria rhythmicity tier (High as default). Each axis is accompanied by a confidence value and the criterion weight of the winning model.

The pairwise comparisons are retained in full, and the amplitude and phase differences, their p-values, and the corresponding deltas are reported for every pair of groups. However, the biological category assigned to each pair now follows the selected model rather than the isolated pairwise test. A pair of rhythmic groups placed in the same block is called unchanged, whereas a pair placed in different blocks is called changed, with the responsible component taken from the pairwise amplitude and phase p-values. This keeps the target-level and the pairwise descriptions consistent while leaving the underlying test visible.

Because the rhythm-difference test is only read for targets with two or more rhythmic groups, correcting it across all targets may be over-conservative. CODAC_multi therefore offers a screened correction, in which the correction family is restricted to the targets passing a corrected pooled shared-rhythm screen. The two tests are orthogonal, so the screen does not bias the rhythm-difference test, and the resulting corrected value is never larger than the genome-wide one.

Taken altogether, CODAC_multi provides a target-level summary of how rhythms are organized across multiple groups, the model of change, and the confidence in it, while retaining the pairwise detail from which that model is built.

## 5 Discussion

Circadian bioinformatic analysis depends heavily on experimental design, including sampling interval and replicate number, and on the specific algorithm applied to the data. Different algorithms rest on different premises, statistical models of rhythmicity, processing steps, waveform assumptions, and criteria for calling a change in rhythm parameters. A recent study [20] showed that these differences in design and method choice produce markedly different results: differential-rhythmicity algorithms applied to the same dataset, using comparable statistical thresholds, identified different sets and numbers of rhythmic transcripts. These discrepancies mostly reflect how each method defines rhythmicity and differential rhythmicity, rather than a simple split between correct and incorrect calls. Experimental variables such as animal background, diet, light schedule, housing, and sample processing add further variability to RNAseq-based circadian studies. This study [21] processed 57 mouse liver time series and found limited overlap in the rhythmic genes identified across datasets; most of this discordance traced back to technical factors, such as library preparation, with biological variables such as age, sex, feeding schedule, lighting, sampling density, and replicate number contributing as well. Notably, the phase of core clock genes remained consistent across all 57 datasets, indicating that the core clock’s temporal architecture is more robust to these sources of variability than downstream transcriptional output.

An important feature of CODAC is that rhythmicity is not defined by a p-value alone. CODAC integrates statistical significance with goodness of fit, amplitude cutoff, and an interval-of-variability criterion to classify each target as low, medium, high, or extremely high confidence. This multiparameter strategy provides a graded assessment of rhythmic evidence and separates strongly supported rhythms from patterns that are statistically significant but poorly fit. Such an approach is likely to be particularly valuable in low-resolution datasets (e.g., 6 hours sampling) and in designs with a single replicate per timepoint. In addition, CODAC has a flexible framework. Parameters such as period, minimum model fit, statistical threshold, interval variability, amplitude stringency, missing-value handling, and minimum rhythmicity confidence are all user-adjustable. CODAC therefore offers a flexible package for studying biological rhythms across a range of experimental designs.

In our benchmarking, the medium-confidence category captured the trade-off between sensitivity and specificity. Retaining these targets increased sensitivity to weak or borderline rhythms, but it also increased the number of rhythmic calls and, with it, the risk of overclassification. In our simulated analyses, including medium-confidence targets produced more rhythmic calls than the number of rhythms actually introduced into the simulated dataset. In the cross-method comparison, retaining this category also reduced agreement between CODAC and established methods. By explicitly accounting for this uncertainty rather than forcing a binary arrhythmic/rhythmic split, CODAC offers a more nuanced view of the underlying biology.

CODAC_compare achieved moderate-to-substantial agreement with established methods for arrhythmic classifications, whereas agreement for rhythmic classifications was lower and more variable, in line with [20]. Concordance was generally strongest with diffCircadian and was also evident with CircaCompare and compareRhythms, while DryR showed the lowest agreement. In part, this is not unexpected as DryR assigns each target to the best-supported model by BIC rather than testing amplitude and phase differences against a significance threshold. Importantly, excluding medium-confidence targets consistently improved agreement across all four comparisons. However, agreement should not be read as accuracy: Cohen’s *κ* measures whether two methods classify the same targets similarly after accounting for chance, not which method is biologically correct.

CODAC_multi extends this framework to more complex designs involving three or more groups. Rather than reporting only pairwise contrasts, CODAC_multi identifies which groups share a common rhythmic pattern and which form distinct rhythmic blocks, comparing all compatible grouping configurations by BIC or AICc and reporting the best-supported model together with a confidence value and information-criterion gap. This lets users see, in a single output, the clustering of targets that behave similarly across the comparisons they define, while the underlying pairwise amplitude and phase statistics remain available to describe the specific components of each difference.

Taken altogether, our findings position CODAC as a flexible addition to the existing toolkit for studying circadian rhythms. By combining fixed and flexible waveform models, pairwise and multi-group comparisons, a graded multiparameter confidence system, and visualization tools such as rhythm heatmaps, CODAC lets users tailor the analysis to their experimental design.

## 6 Statements and Declarations

### Funding

Leonardo Vinicius Monteiro de Assis is supported by the Knut and Alice Wallenberg Foundation as a Wallenberg Molecular Medicine Fellow, the Jeanssons Foundation, and the German Research Foundation (Deutsche Forschungsgemeinschaft) through grant 541063275—TRR B04. Thiago Parente da Silveira is supported by CNPq (grant 407147/2023-3) and FAPESP (grant 2023/08706-1).

### Competing Interests

The authors declare that they have no competing interests.

### Author Contributions

**Conceptualization** : TPS and LVMA.

**Methodology** : TPS and LVMA.

**Validation** : KL and TN.

**Data Curation** : KL and TN.

**Writing—Original Draft** : TPS and LVMA.

**Writing—Review and Editing** : All authors.

### Data Availability

No new datasets were generated in this study. Publicly available datasets used in the analyses are identified by their corresponding accession numbers in the Methods section. All data used for all figures are available in Table S1.

### Code Availability

CODAC is open-source and freely available at https://github.com/ThiagoPSilveira/CODAC. The repository contains the source code for CODAC_single, CODAC_flex, CODAC_compare, and CODAC_multi, together with documentation and tutorials for installation and use.

### Artificial Intelligence Statement

CODAC was designed with Julius and was subsequently tested with ChatGPT and Gemini. The final R package was completed using Claude. Grammarly was used to improve the clarity of the manuscript. The authors take full responsibility for the accuracy, integrity, and reproducibility of the final software and manuscript.

## Notes

### Competing Interest Statement

The authors have declared no competing interest.

## References

[1] Takahashi, J. S. Transcriptional architecture of the mammalian circadian clock. Nature Reviews Genetics 18, 164–179 (2017).

[2] de Assis, L. V. M. & Oster, H. The circadian clock and metabolic homeostasis: Entangled networks. Cellular and Molecular Life Sciences 78, 4563–4587 (2021).

[3] Koike, N. et al. Transcriptional architecture and chromatin landscape of the core circadian clock in mammals. Science 338, 349–354 (2012).

[4] Mauvoisin, D. Circadian rhythms and proteomics: It’s all about posttranslational modifications! Wiley Interdisciplinary Reviews: Systems Biology and Medicine 11, e1450 (2019).

[5] Menet, J. S., Rodriguez, J., Abruzzi, K. C. & Rosbash, M. Nascent-seq reveals novel features of mouse circadian transcriptional regulation. eLife 1, e00011 (2012).

[6] Robles, M. S., Cox, J. & Mann, M. In-vivo quantitative proteomics reveals a key contribution of post-transcriptional mechanisms to the circadian regulation of liver metabolism. PLoS Genetics 10, e1004047 (2014).

[7] Hughes, M. E., Hogenesch, J. B. & Kornacker, K. JTK_CYCLE: An efficient nonparametric algorithm for detecting rhythmic components in genome-scale data sets. Journal of Biological Rhythms 25, 372–380 (2010).

[8] Thaben, P. F. & Westermark, P. O. Detecting rhythms in time series with RAIN. Journal of Biological Rhythms 29, 391–400 (2014).

[9] Parsons, R., Parsons, R., Garner, N., Oster, H. & Rawashdeh, O. CircaCompare: A method to estimate and statistically support differences in mesor, amplitude and phase, between circadian rhythms. Bioinformatics 36, 1208–1212 (2020).

[10] Pelikan, A., Herzel, H., Kramer, A. & Ananthasubramaniam, B. Venn diagram analysis overestimates the extent of circadian rhythm reprogramming. The FEBS Journal 289, 6605–6621 (2022).

[11] Weger, B. D. et al. Systematic analysis of differential rhythmic liver gene expression mediated by the circadian clock and feeding rhythms. Proceedings of the National Academy of Sciences 118, e2015803118 (2021).

[12] Hughes, M. E. et al. Guidelines for genome-scale analysis of biological rhythms. Journal of Biological Rhythms 32, 380–393 (2017).

[13] Parsons, R., Jayasinghe, O., White, N., Chunduri, P. & Rawashdeh, O. GLMMcosinor: Flexible cosinor modeling to characterize rhythmic time series using a generalized linear mixed modeling framework. bioRxiv (2025). Preprint.

[14] de Assis, L. V. M., Demir, M. & Oster, H. Nonalcoholic steatohepatitis disrupts diurnal liver transcriptome rhythms in mice. Cellular and Molecular Gastroenterology and Hepatology 16, 341–354 (2023).

[15] de Assis, L. V. M. et al. Hepatocyte circadian clocks control cholesterol metabolism and protect from metabolic dysfunction-associated steatohepatitis (MASH). Cellular and Molecular Gastroenterology and Hepatology 0 (2026).

[16] Zhang, R., Lahens, N. F., Ballance, H. I., Hughes, M. E. & Hogenesch, J. B. A circadian gene expression atlas in mammals: Implications for biology and medicine. Proceedings of the National Academy of Sciences of the United States of America 111, 16219–16224 (2014).

[17] de Assis, L. V. M. et al. Thyroid hormone receptor beta (THRB) dependent regulation of diurnal hepatic lipid metabolism in adult male mice. npj Metabolic Health and Disease 2, 1–11 (2024).

[18] Deota, S. et al. Diurnal transcriptome landscape of a multi-tissue response to time-restricted feeding in mammals. Cell Metabolism 35, 150–165.e4 (2023).

[19] Rubio-Ponce, A. et al. Combined statistical modeling enables accurate mining of circadian transcription. NAR genomics and bioinformatics 3, lqab031 (2021).

[20] Miao, L., Weidemann, D. E., Ngo, K., Unruh, B. A. & Kojima, S. A comparative study of algorithms detecting differential rhythmicity in transcriptomic data. Bioinformatics and Biology Insights 18, 11779322241281188 (2024).

[21] Brooks, T. G., Manjrekar, A., Mrčela, A. & Grant, G. R. Meta-analysis of diurnal transcriptomics in mouse liver reveals low repeatability of rhythm analyses. Journal of Biological Rhythms 38, 556–570 (2023).

